# Fine-scale behaviour and population estimates suggest low exposure but do not exclude high sensitivity to bycatch for Endangered sooty albatrosses

**DOI:** 10.1101/2024.04.14.589402

**Authors:** Aymeric Fromant, Julien Collet, Cécile Vansteenberghe, Raphäel Musseau, Dominique Filippi, Karine Delord, Christophe Barbraud

## Abstract

Recent developments in assessing species-specific seabird bycatch risks demonstrated that fine-scale approaches are essential tools to quantify interactions with fishing vessels and understand attraction and attendance behaviours. Matching boats movements with birds tracking data specifically allows to investigate seabird-fishery interaction for cryptic species for which on-board information is critically lacking. The sooty albatross (*Phoebetria fusca*) overlaps with fisheries throughout its range and is known to be vulnerable to incidental bycatch. Combining GPS and behaviour data from individuals from Crozet Islands and boat locations during the incubation period, we investigated interactions of sooty albatrosses with fisheries in the southern Indian Ocean. Individuals foraged mostly in sub-tropical international waters, where they only encountered a small number of boats. The low interaction rate during this period may suggests that sooty albatrosses are not strongly attracted towards fishing vessels. However, this result should be interpreted with caution due to the low sample size and fishing effort during the study period, as these observations may conceal a higher bycatch risk during intense fishing effort and/or energetically demanding periods. The species conservation status requires further data to be collected throughout the annual cycle to provide an accurate assessment of the threat.

## Introduction

Due to the worldwide impacts of climate change and anthropic activities on oceanic ecosystems, seabirds are one of the most threatened groups of birds (Croxall et al. 2012, Dias et al. 2019). Facing threats both on land and at sea, more than half of all seabird species have declining trends, and a third of all species are globally threatened (BirdLife International 2022). Incidental bycatch (hereafter bycatch) from fisheries has clearly been identified to be one of the major threats to seabirds at sea, with large-scale longline fisheries having the greatest impact in terms of severity and scope (Dias et al. 2019). Despite the improvement in implementation of mitigation regulations over the last decades, seabirds still suffer from substantial bycatch risks (Votier et al. 2023).

Effective mitigation measures have been implemented for longline vessels, such as discard management (Bull 2007), hook-shielding devices (Sullivan et al. 2018), night setting (Brothers et al. 1999), bird-scaring lines (Domingo et al. 2017) and weighted branch lines (ACAP 2017, Paterson et al. 2017). Yet, the absence of population recovery for some species indicates that the implementation of mitigation measures remains relatively inadequate (Pardo et al. 2017). Our understanding of interaction processes is particularly limited for cryptic species for which the lack of knowledge on bycatch rates and interaction behaviour is scarce. Obtaining detailed on-board information can be challenging owing to low coverage of vessels by official observers among fisheries (Winnard et al. 2018). Therefore, increasing research effort to better understand the species behaviour related to industrial fishing activities is needed to adapt conservation measures and their effective compliance.

Until recently, bycatch risk was traditionally assessed through the coarse overlap estimates between fishing effort and seabird foraging area over large spatial and temporal scales (Clay et al. 2019, Heerah et al. 2019). Although this approach provides useful information to infer risk assessments at the population level and through the life cycle of a species, the absence of fine-scale data limits our understanding of when and how seabirds are exposed to bycatch. In particular, this method does not discriminate actual overlap versus interaction rate (i.e. theoretical vs real risk), which may mislead in assessing threats and subsequent mitigation measures. The recent possibility to access publically available Automatic Identification System (AIS), providing the locations and types of all declared vessels, quickly became the most suited solution to obtain accurate information on the actual time seabirds spend interacting with fishing boats (Winnard et al. 2018). This new approach appears to be especially relevant to provide fine-scale bycatch risk for species ranging outside the national Exclusive Economic Zones (EEZ), where mitigation measures are particularly difficult to implement (Dias et al. 2019). In addition, deploying radar-detecting devices on seabirds was shown to be a necessary complementary approach to detect any vessels not using AIS (Weimerskirch et al. 2018a), such as illegal and unregulated fishing boats. These fine-scale approaches were successfully applied to investigate bycatch risks for several seabird species (Weimerskirch et al. 2020, Corbeau et al. 2021a), providing crucial information on their interaction behaviour towards fishing vessels, and helping to assess the proportion of illegal vessels encountered. However, while these studies focused on relatively abundant and conspicuous species, such as the wandering albatross (*Diomedea exulans*), that are strongly attracted to fishing vessels, knowledge of more cryptic species has remained limited.

The sooty albatross (*Phoebetria fusca*, endangered species IUCN) was recently classified as one of the procellariform species to be most exposed to bycatch risks (Reid et al. 2023), confirming the potential contribution of fisheries bycatch mortality to the observed multi-decade population decrease throughout the Southern Ocean (Cuthbert & Sommer 2004, Delord et al. 2008, Rolland et al. 2010, Schoombie et al. 2016, Weimerskirch et al. 2018b). Nonetheless, the sooty albatross is also described as a species interacting only rarely with fishing vessels (Griffiths 1982, SIOFA 2024, unpublished French Southern Breeding Seabird Survey database), emphasising the need to quantify fine-scale seabird-fishery interactions for such species. Using AIS, Banda et al. (2023) demonstrated that sooty albatrosses from Marion Island displayed much lower exposure and attraction to fishing vessels compared to other species in the Southern Ocean, such as the wandering albatross (Corbeau et al. 2021a, Carneiro et al. 2022) or the white-chinned petrel (*Procellaria aequinoctialis*) (Banda et al. 2023). However, the apparent discrepancy between the sooty albatross behaviour and its exposure to bycatch fades in when considering the combined effects of the small population and the life history of the species. In particular, sooty albatrosses are biennial breeders with a single egg clutch, meaning that populations from this species rely on a long-term high survival rate to maintain a stable, small population. In this context, any bycatch event, even with a low exposure risk, may have a significant impact at the population level (Dillingham & Fletcher 2011).

In addition, variations in foraging habitats depending on the geographical location of populations can also lead to differences in overlap and attraction to fishing vessels (Soriano-Redondo et al. 2016, Cianchetti-Benedetti et al. 2018). For example, wandering albatrosses from Crozet Islands appear to attend boats for shorter duration than conspecifics from Kerguelen Islands, but encountered more illegal fishing vessels due to their specific foraging habitat (Corbeau et al. 2021b). These results exemplify the need to develop bycatch risk assessments for sooty albatrosses at the population level, which also must include information about exposure to illegal fisheries.

The Crozet Islands used to host one of the largest population of sooty albatrosses in the southern Indian Ocean, although the number of breeding pairs has declined by more than 81 % since 1980 (Fig. 1; Delord et al. 2008, Weimerskirch et al. 2018b). Therefore, it is crucial to assess the bycatch risk that this population is exposed to, which implies quantifying and understanding interactions at fine spatial and temporal scales. Combining seabird GPS tracking data, radar detector and vessel AIS data, this study aimed to 1) obtain accurate information (occurrence and location) on interactions between fisheries (including illegal fishing vessels) and sooty albatrosses from Crozet Islands during the incubation period; and 2) investigate the behaviour of individuals encountering and attending boats (fishing vessels or not) by determining if the presence of a vessel influences their searching activity.

**Fig. 1:**
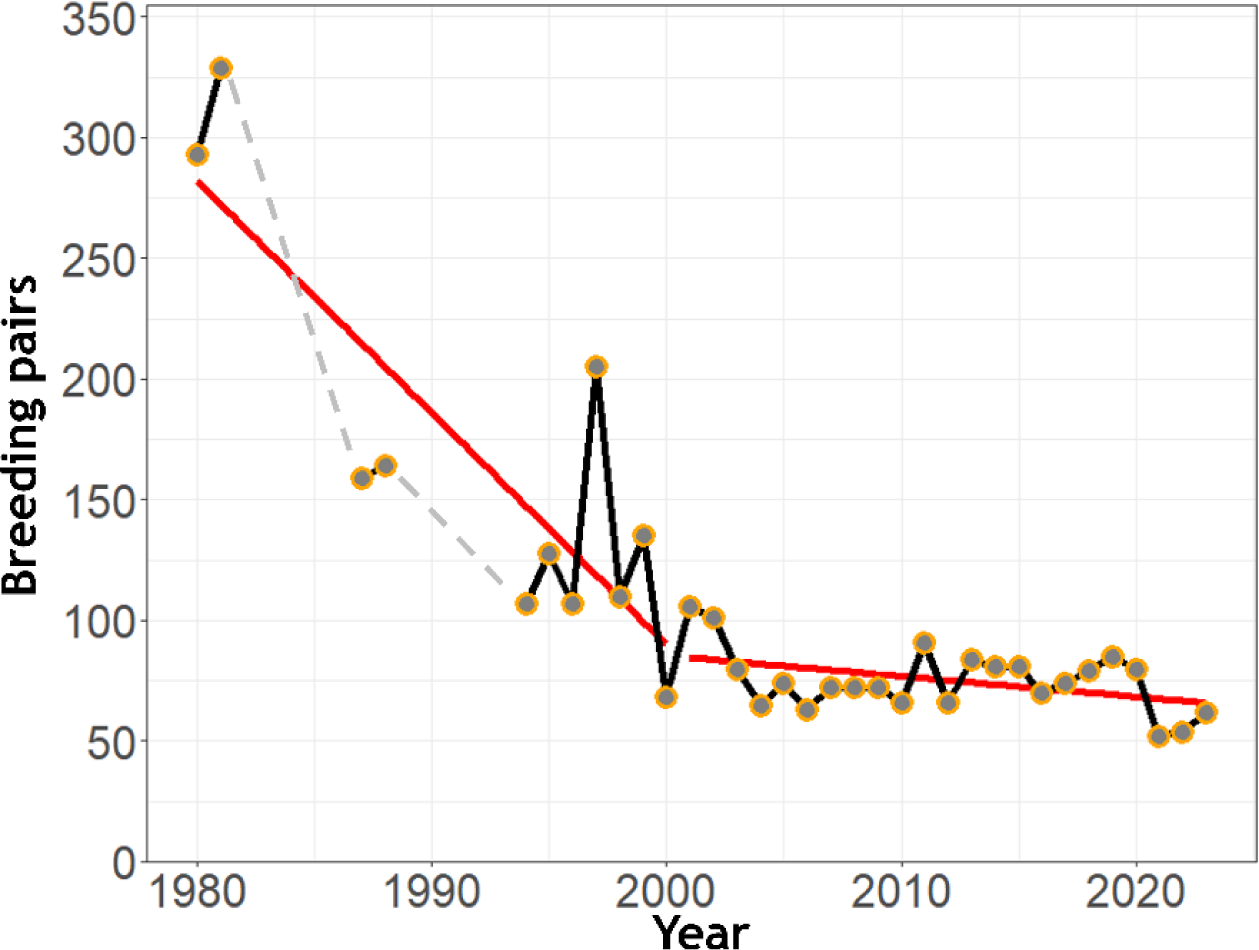
Annual number of breeding pairs of sooty albatrosses at Ile de la Possession, Crozet archipelago. Orange dots correspond to annual ground counts of breeding pairs. The red lines indicate linear trends for the periods 1980-2000 and 2001-2023.

## Methods

### Study site and study species

The study was carried out at Ile de la Possession (Crozet Islands, southern Indian Ocean; 46°S 51°E) between November and December 2022. Crozet archipelago, situated between the Sub-Antarctic and the Antarctic Polar fronts, used to host nearly 16% of the world’s population of sooty albatrosses (Delord et al. 2008, 2013). The population of sooty albatrosses on the archipelago (regrouping five main islands) was last estimated by Jouventin et al. (1984), with 2,335 breeding pairs. Since, the population of Ile de la Possession was used as a reference to estimate Crozet archipelago’s population trend (Delord et al. 2008). This population has undergone an important decline between 1981 and 2000 at an average rate of 4.2% per year, followed by a slower decline of 1.9% per year over the last two decades (Fig. 1). With a 81% decrease between 1980-81 and 2022-23, the whole Crozet population is estimated to have reduced down to 444 annual breeding pairs.

Sooty albatross is a biennial breeder. Individuals are present at Crozet between late August to late June, incubating a single egg from early October to mid-December with chicks fledging late May (Weimerskirch et al. 1986). During the incubation period, both male and female alternate incubation shifts with foraging trips lasting on average 11 days (Weimerskirch et al. 1986).

### Tracking data and fishing effort

A total of 13 adult sooty albatrosses were equipped with XAIS-Sputnik loggers developed by Sextant Technology (recording location, wet/dry information, and equipped with a radar detector) during the incubation period in order to record their at-sea movements and behaviour. The devices were programmed to record locations every 10 min for at least one trip in late incubation. Individuals were captured on the nest at the estimated end of their incubation shift, and loggers were attached to the back feathers using waterproof tape (Tesa 4651, Beiersdorf AG, Germany). Although in the present study the equipped individuals were not weighed to limit the duration of handling, the total mass of logger attachments (52 g) was estimated to correspond to 2.1 % of body mass (average body mass = 2.5 kg; Marchant & Higgins 1990). Handling duration was 8.6 ± 1.4 min for deployment, and 4.4 ± 2.5 min for retrieval.

Automatic identification systems of vessels operating in the study area were downloaded from Global Fishing Watch (https://globalfishingwatch.org) using the R package *gfwr* (Merten et al. 2016). The Global Fishing Watch website provides tracking data from available AIS and combines information acquired through vessel monitoring systems that are made available through partnerships with governments. For each vessel, location, identification name, type of vessel (fishing or not), and activity were obtained for the same temporal and spatial extent as the bird tracked. GPS loggers deployed on sooty albatrosses were combined with a radar detector. This system was developed to detect vessels that do not use AIS, in particular illegal fishing vessels. This method allows the detection of vessels with radar emitting within a 5 km range from equipped birds (Weimerskirch et al. 2018a). Therefore, vessels illegally deactivating their AIS will only be recorded if there is a close encounter.

Fishing effort from longline fisheries was provided by the Southern Indian Ocean Fisheries Agreement (SIOFA; sub-areas 1, 2, 3a and 3b) and the Commission for the Conservation of Antarctic Living Resources (CCAMLR; sub-areas 58.6 and 58.7). The number of hooks deployed was pooled by month and sub-areas to determine the spatio-temporal variation of the fishing effort in the area used by sooty albatrosses from Ile de la Possession.

### Behavioural and data analysis

For tracking data of sooty albatrosses, all on-land locations were removed from GPS tracks, and locations were interpolated with a time step of 10 min to correct for any unequal sampling frequencies. For each complete foraging trip, the following basic parameters were determined: trip duration (h), total horizontal distance travelled (km) and maximum distance from the colony (km). Sooty albatross locations and AIS data were then spatio-temporally matched following Weimerskirch et al. (2020) resulting in a dataset with, for each bird location, information about the nearest vessel transmitting AIS. Distance thresholds of 100 km (boat seascape), 30 km (boat encountered) and 5 km (boat attended) were used to classify and determine the behaviour of sooty albatrosses when foraging within each of these categories. Within the 100 km range, the boat was considered to be available in the seascape of the individual, while the 30 km range corresponds to the maximum distance at which an albatross can visually detect a vessel (Collet et al. 2015). The 5 km threshold relates to the distance at which seabirds perform specific foraging behaviours when reaching this proximity of a vessel (Collet et al. 2015, Corbeau et al. 2019, Weimerskirch et al. 2020).

The expectation maximization binary clustering (EMBC) was used to infer the at-sea foraging behaviour of sooty albatrosses using the R package *EMbC* (Garriga et al. 2016). Using travel speed and turning angle between subsequent locations, this method classifies the movement of seabirds into four different categories: travelling-commuting (high speed, low turn), extensive searching (high speed, high turn), intensive searching (low speed, high turn) and resting on the water (low speed, low turn). This method is well suited to interpreting ecologically meaningful behaviours for procellariform species, including albatrosses (De Grissac et al. 2017).

In addition, behavioural information was collected using wet and dry data. When submerged, the loggers recorded the number of “wet” every second for a duration of 2 min, restarting the process again if the device was still submerged after 2 min. This method allowed to save battery and memory space while recording high-resolution data. The dataset was merged with the bird location file to obtain a percentage of time spent submerged per location. This information was then combined with the EMBC classification to refine the interpretation of sooty albatross foraging behaviour.

All processing and statistical analyses were conducted in the R statistical environment (R Development Core Team 2022). For each trip, the number of vessels (fishing or not) in seascape, encountered or attended was determined. The duration during which individuals stayed within 100 km or 30 km for the events that ended up with an encounter, or attendance, or neither, were compared using Mann-Whitney *U*-tests (non-parametric). Statistical comparison of the four behavioural classes between the different radius distances (100, 30 and 5 km) was impaired because of the small number of boat attendances (four instances).

## Results

### Foraging trips

A total of 12 devices were retrieved out of the 13 deployed on breeding sooty albatrosses (recovery rate = 92.3%). The last device was never retrieved due to breeding failure. In addition, no data was recovered from one device, likely due to water logging. Successful devices collected tracking data for 11 trips at the end of the incubation period (11 individuals; between 27-Nov and 16-Dec). For two individuals, an additional three trips were recorded during early chick-rearing (between 19-Dec and 25-Dec) (Table 1). From the 11 individuals tracked (14 foraging trips), a total of 2,682 hours of tracking were recorded, corresponding to 112 days at sea.

**Table 1:** Boats in seascape, encountered and attended by sooty albatrosses from Ile de la Possession. Each row corresponds to one trip, indicating the number of instances this individual entered within 100, 30 or 5 km of a vessel, and the total duration (h) during which the individual was within this distance during the whole trip. All the foraging trips were during late incubation, except trips SA-111 (2), SA-114 (2) and SA-114 (3) that were during early brooding stage (see Fig. 3).

| Individual<br>(Trip) | Trip duration<br>(h) | < 100 km |  | < 30 km |  | < 5 km |  |
| --- | --- | --- | --- | --- | --- | --- | --- |
|  |  | Instances<br>(fishing vessel) | Duration (h)<br>(fishing vessel) | Instances<br>(fishing vessel) | Duration (h)<br>(fishing vessel) | Instances<br>(fishing vessel) | Duration (h)<br>(fishing vessel) |
| SA-105 | 249 | 10<br>(1) | 112.7<br>(1.7) | 4 | 14.0 | 2 | 1.3 |
| SA-108 | 316 | 5<br>(3) | 74.0<br>(36.3) | 3<br>(2) | 10.7<br>(7.6) | 1<br>(1) | 0.7<br>(0.7) |
| SA-101 | 260 | 5<br>(1) | 45.3<br>(3.7) | 3 | 8.0 | 1 | 0.7 |
| SA-106 | 243 | 6<br>(2) | 56.0<br>(29.0) | 2<br>(1) | 5.0<br>(1.7) | - | - |
| SA-111 (1) | 235 | 4<br>(2) | 42.7<br>(19.4) | 2<br>(1) | 3.7<br>(2.7) | - | - |
| SA-111 (2) | 62 | 2<br>(1) | 28.7<br>(13.7) | 2<br>(1) | 6.7<br>(1.7) | - | - |
| SA-107 | 260 | 3<br>(1) | 25.0<br>(4.0) | - | - | - | - |
| SA-104 | 237 | 4 | 42.7 | - | - | - | - |
| SA-110 | 213 | 2<br>(1) | 8.3<br>(5.3) | - | - | - | - |
| SA-114 (2) | 78 | 2<br>(1) | 10.0<br>(4.7) | - | - | - | - |
| SA-111 (3) | 17 | 1<br>(1) | 6.0<br>(6.0) | - | - | - | - |
| SA-112 | 157 | - | - | - | - | - | - |
| SA-114 (1) | 218 | - | - | - | - | - | - |
| SA-103 | 136 | - | - | - | - | - | - |

Adult sooty albatrosses travelled on average 5285 ± 1320 km per foraging trip during incubation, for a mean duration of 229 ± 49 hours at sea, and a mean maximum distance from the colony of 1340 ± 340 km. The three trips in early chick-rearing were much shorter, with a mean travelled distance of 908 ± 434 km, a mean duration of 52 ± 31 hours and a mean maximum distance from the colony of 442 ± 30 km. During incubation, all individuals travelled further than the extent of the EEZ of Crozet Islands, foraging mostly north of the Sub-Antarctic (SAF) and Sub-tropical fronts (STF) (Fig. 2). Sooty albatross allocated proportionally more time searching (intensive and extensive searching) when they were in sub-tropical open waters (25.0%) compared to when foraging in sub-antarctic waters (16.8%).

**Fig. 2:**
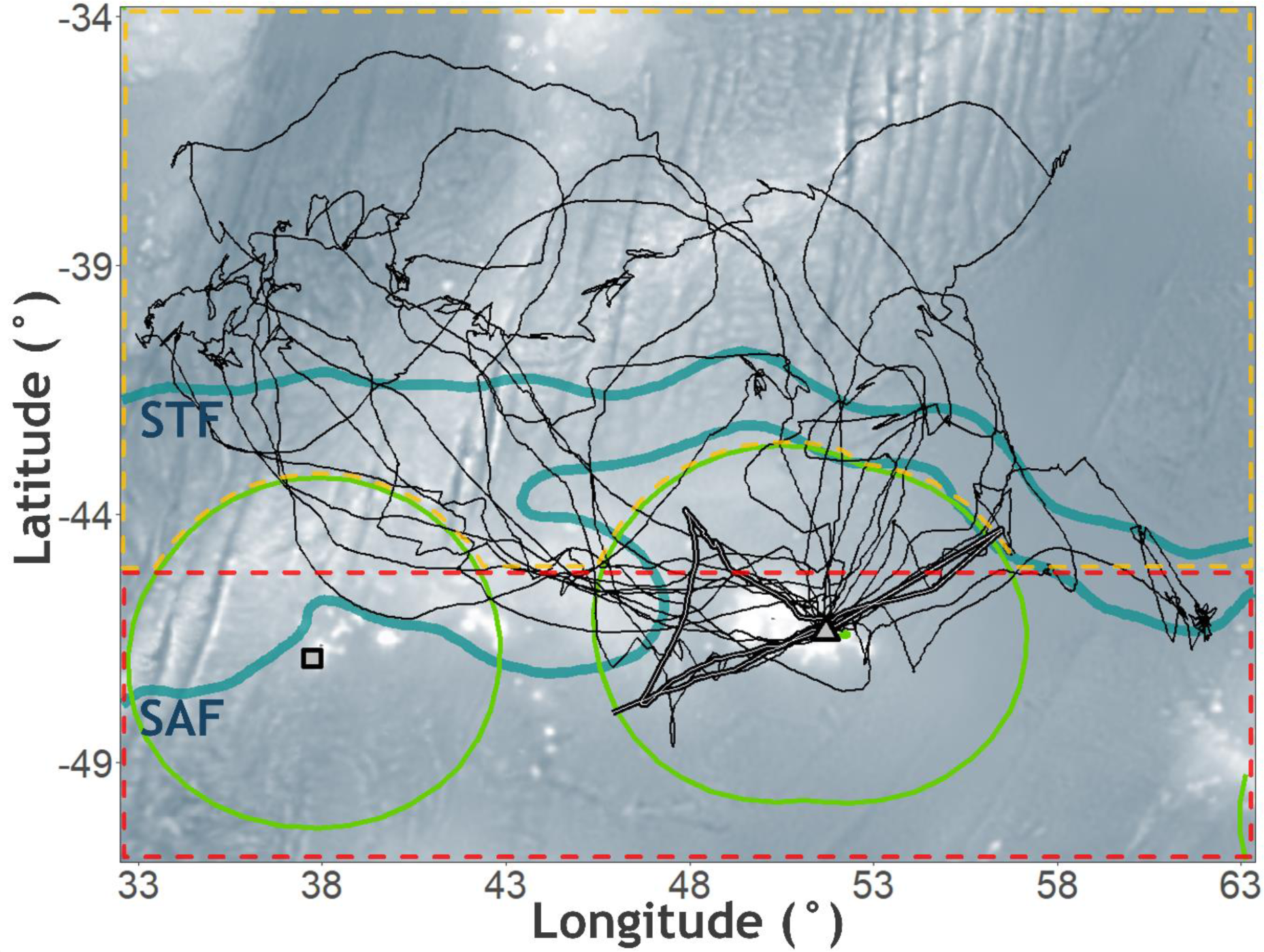
Foraging trips of sooty albatrosses from Ile de la Possession, Crozet Islands. The thin black tracks correspond to foraging trips during the incubation period, while the thick black and white tracks (within the Crozet EEZ) show the three trips in early brooding stage. STF = Subtropical Front; SAF = Sub-Antarctic Front (Kim & Orsi 2014). The circular green shapes correspond to the Exclusive Economic Zones (EEZs) of Crozet Islands (grey triangle) and Marion and Prince Edward Islands (grey square). The dotted lines correspond to the SIOFA (orange) and the CAMMLR (red) areas.

### Boatscape description

During the period when sooty albatrosses were tracked (27-Nov to 25-Dec), 310 vessels were identified (using AIS data set) in the study area, including 26 fishing vessels (8.4%). The great majority of ship activity was concentrated in a large area north of −39°, while fishing activity was mostly restricted south of this zone (Supplementary Fig. 1). Sooty albatrosses were recorded 44 times within 100 km of a vessel, of which 14 corresponded to a fishing vessel (31.8%) (Table 1). Of these instances, individuals were recorded 16 times (six different individuals) within 30 km of a vessel (five fishing vessels; 31.3%), and four of these encounters (3 different individuals) ended up with an individual recorded within 5 km of a vessel (one fishing vessel; 25%) (Fig. 3; Table 1). For three of the 14 trips, the individuals were never recorded within 100 km of a vessel (Table 1).

**Fig. 3:**
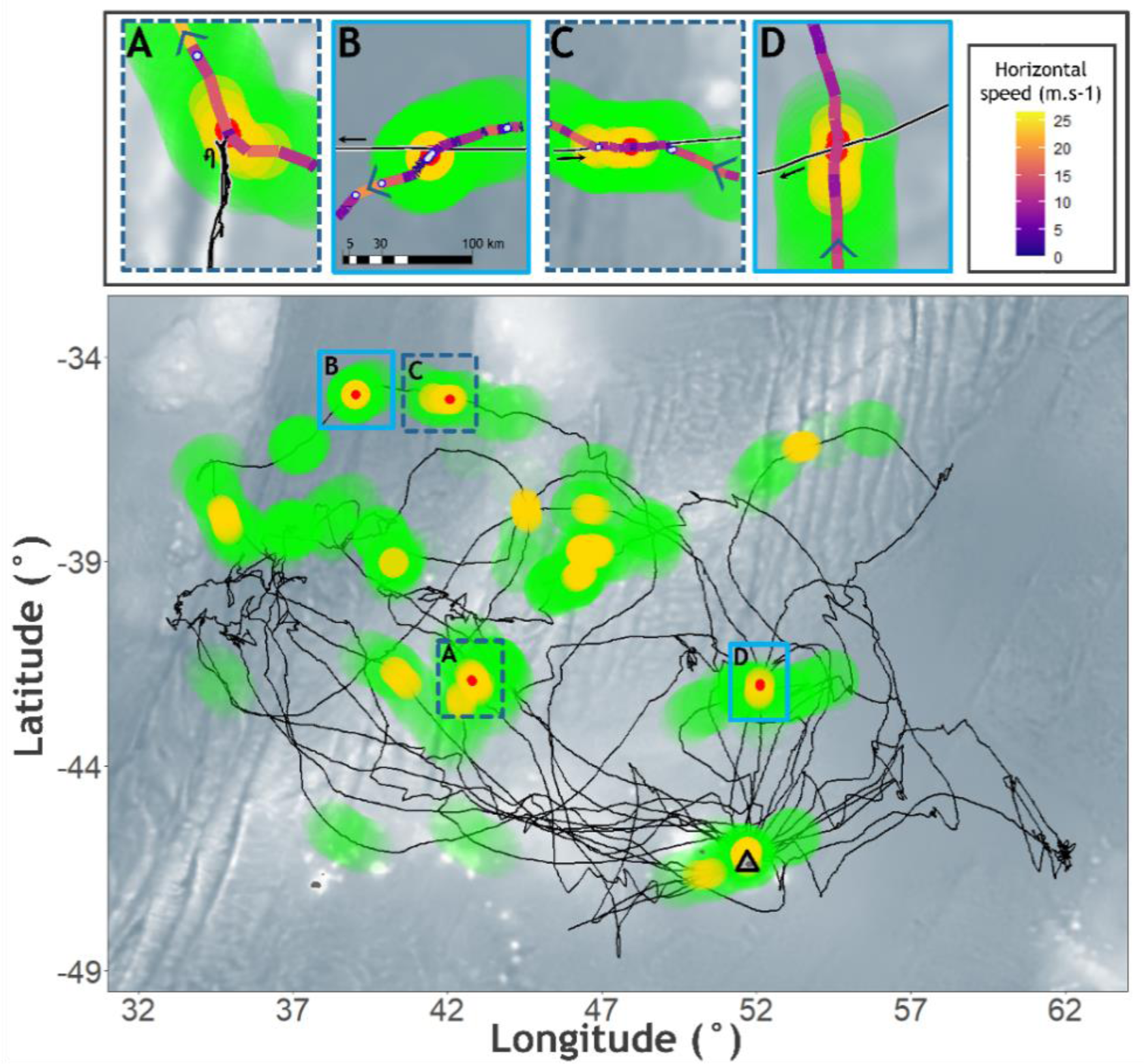
Position of vessels within seascape (< 100 km; green dots), vessels encountered (< 30 km; yellow dots), and vessels attended (< 5km; red dots) by sooty albatrosses (black track) from Ile de la Possession. The four cases for which an individual was within 5 km of a vessel are enlarged on the top of the caption (A to C). For each case, arrows indicate the direction of the individual and the vessel, the colour of the bird track corresponds to its speed, and the white dots show events when the device was submerged (note that the green, yellow and red dots are representational only and are not to scale). Cases A and C (doted dark blue) corresponds to the instances when the radar detection device detected the vessels. Case A was the only instance for which a sooty albatross was within 5 km of a fishing vessel.

Two vessels were detected using radar detection devices deployed in combination with tracking devices attached on the birds. They both matched a vessel attendance (individual within 5 km) detected with the vessel AIS (Fig. 3). One of these was a fishing vessel. There was no detection of any vessel that was not identified within the Global Fishing Watch AIS data base.

For each event of sooty albatrosses being within 100 km of a vessel, individuals were within this zone on average for 4.9 ± 3.5 hours (n = 44). They remained 1.7 ± 1.2 hours within 30 km (n = 16), and 20 ± 0 min (n = 4) within 5 km of a vessel. There were 7 cases in which individuals never went within 30 km of a vessel (6 different individuals). In these instances, sooty albatrosses stayed significantly less time within 100 km compared to the cases for which individuals were found within 30 km of a vessel (3.9 ± 3.2 vs 6.7 ± 3.5 hours; Mann-Whitney *U* test, *U* = 350, *P* = 0.002). However, there was no difference in the duration spent within 100 km of a vessel between cases with individuals recorded within 30 km but never entered within 5 km, and cases when individuals were found within 5 km of a vessel (6.9 ± 3.5 vs 5.9 ± 3.7 hours; Mann-Whitney *U* test, *U* = 18.5, *P* = 0.543). Similarly, there was no difference between these two groups for the duration spent within 30 km of a vessel (1.5 ± 1.3 vs 1.7 ± 0.9 hours; Mann-Whitney *U* test, *U* = 29.5, *P* = 0.540).

### Sooty albatross foraging behaviour and boat encounters

When within 100 km, 30 km, or 5 km, sooty albatrosses did not present any clear change in behaviour. The EMBC model appeared to indicate no clear difference in the proportion of time searching when individuals were within 100 km, 30 km, or 5 km of a vessel compared to when farther away from a vessel (Fig. 4). Similarly, there was no clear pattern in time spent wet (proxy of feeding event; Fig. 4), nor in term of speed or if individuals aimed towards or followed the vessels during encounters (within 30 km) (Fig. 3A-D; Fig. 5). In the four instances for which the birds were within 5 km of a boat, none of the individuals stopped (no wet data), and all of them remained no more than 20 min within this distance of the vessel.

**Fig. 4:**
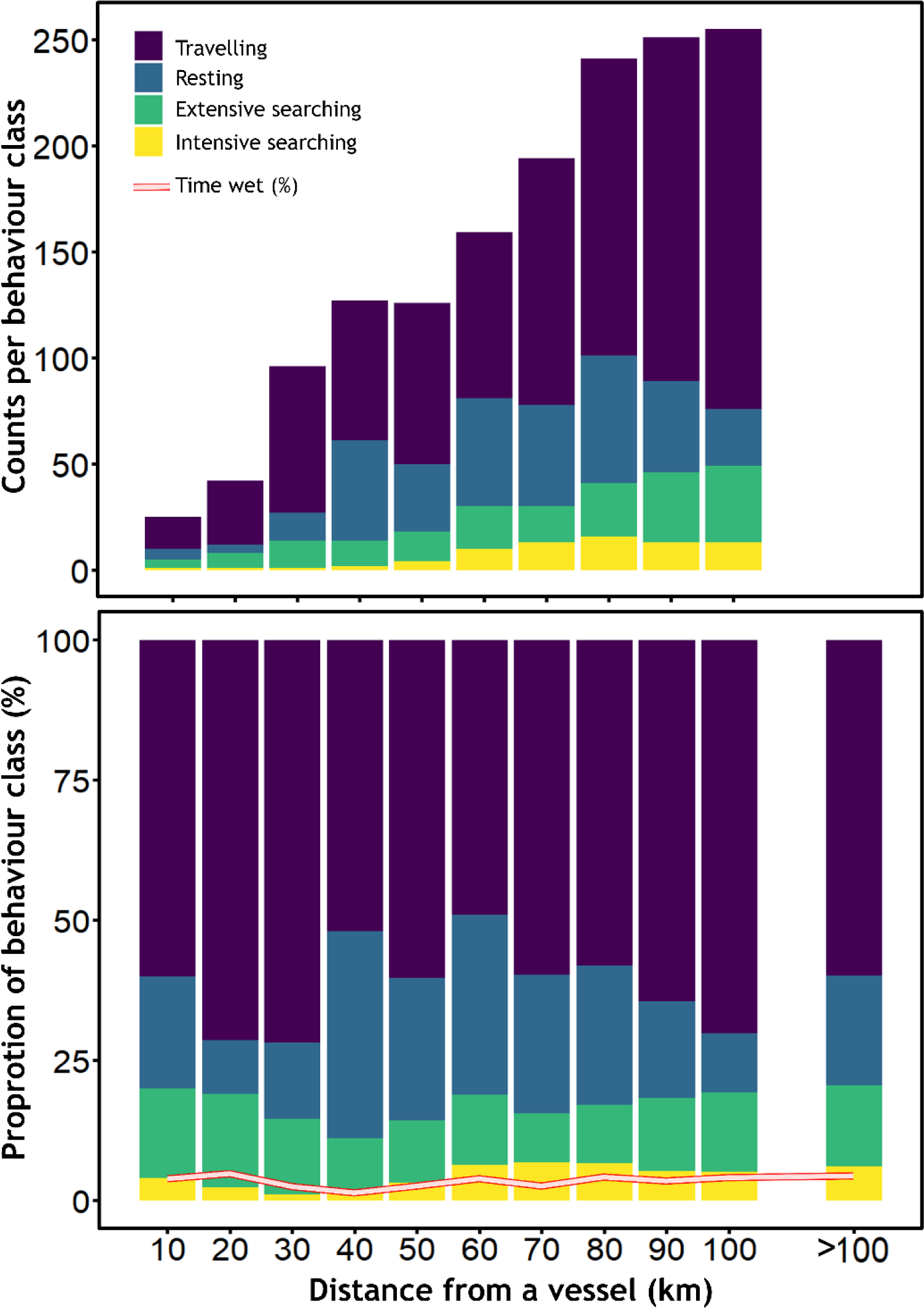
Counts (upper panel) and proportion (lower panel) of behaviour of sooty albatrosses determined using an EMBC model and proportion of time the device was submerged (red and white line). Travelling = high speed, low turn; resting = low speed, low turn; extensive searching = high speed, high turn; intensive searching = low speed, high turn. In the upper panel, counts are not provided for locations >100 km because of the large number of data (>14,000) compared to locations within 100 km.

**Fig. 5:**
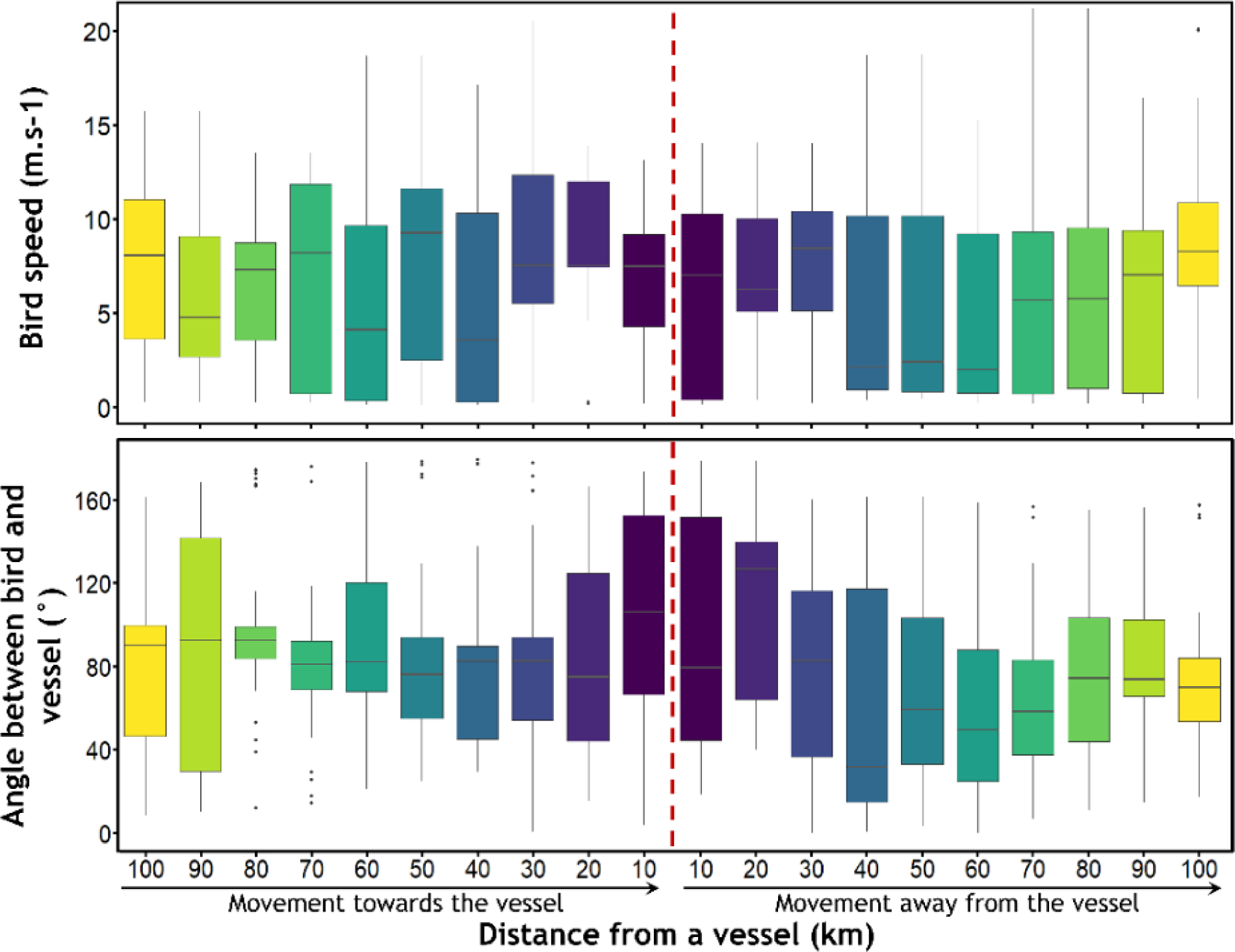
Bearing of sooty albatrosses towards vessels. In the lower panel, an angle of 0° indicates that the individuals were heading towards the vessel, while 180°C indicates that they were heading in the opposite direction.

## Discussion

The present study provides detailed data about the at-sea distribution of sooty albatrosses during the incubation period, quantifying and mapping boat encounters, and contributing to a better understanding of their attraction behaviour towards vessels. The low encounter and interaction rates with fishing vessels during this period may suggest that sooty albatrosses are not strongly attracted towards these boats, and therefore might not be exposed to a high risk of bycatch. However, this low rate of interaction should not dismiss the substantial impacts that a low number of bycatch can generate on small populations, such as the one in the Crozet Archipelago. In addition, the lack of data during periods of high fishing effort, and/or of high energetic demand for individuals, demonstrates the importance of fully assessing bycatch risk throughout the whole annual cycle.

### At-sea distribution and exposure to fishing vessels

At-sea distribution data of breeding albatrosses is crucial for their effective conservation, as these species forage over extensive areas comprising a mix of multiple countries EEZ and High Seas (Thiebot et al. 2014). During the incubation period, sooty albatrosses from Ile de la Possession foraged mostly in open oceanic waters north of the Sub-tropical Front, comparably to individuals from Marion Island (Schoombie et al. 2017, Banda et al. 2023). In this area, sooty albatrosses from both populations overlap with longline and trawling fisheries that have previously documented seabird bycatch (including sooty albatrosses from Ile de la Possession; unpublished French Southern Breeding Seabird Survey database). However, in the present study, and similarly to Banda et al. (2023), there were limited numbers of fishing vessels present in the seascape of the birds, ultimately leading to few encounters. The austral summer is the period during which the fishing effort is the lowest in this area, with eight times fewer hooks deployed in December compared to August (Fig. 6). Because albatrosses cannot detect a vessel presence at a distance higher than 30 km (Collet et al. 2015, Pirotta et al 2018), it is the proportion of fishing boats within the seascape (100 km) that determines the probability leading to potential encounters. The low probability of encounter in December is exemplified by the small proportion of individuals that were within 30 km of a fishing vessel (27% in the present study, 20% in Banda et al. 2023).

**Fig. 6:**
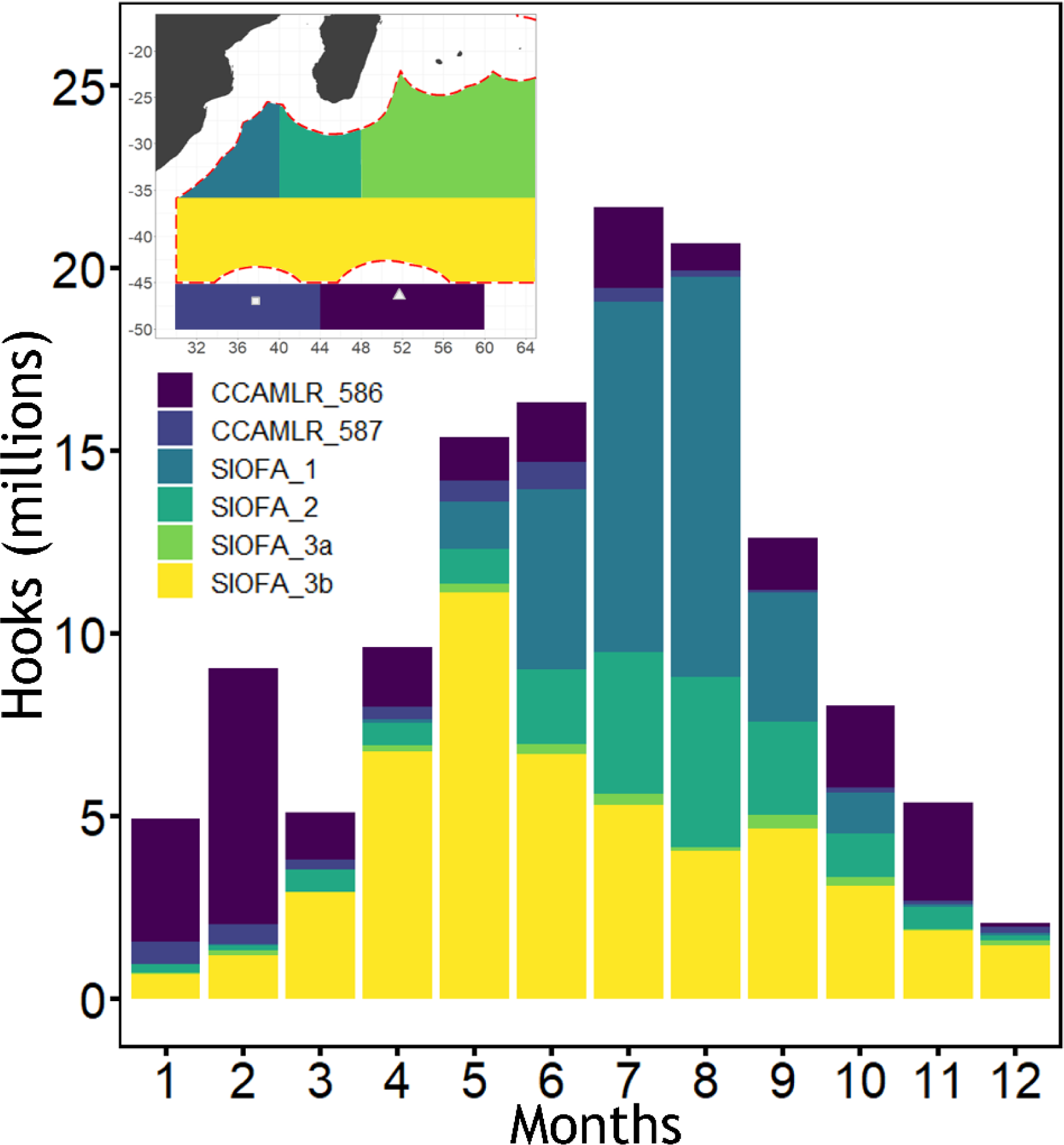
Monthly fishing effort in the areas potentially used by sooty albatrosses from Crozet and Marion islands. Each colour corresponds to the SIOFA (sub-areas 1, 2, 3a and 3b) and the CAMMLR (sub-areas 58.6 and 58.7) areas. The monthly fishing effort was, averaged over five years (2017-2022).

Nonetheless, the perceived low number of interactions with fishing vessels at the individual level hides a much greater bycatch risk when considering the whole population. Based on the actual estimate of 444 breeding pairs for the Crozet Island population, extrapolating the probability of interaction (0.09 per day and individual) would suggest that between two and three individuals are likely to be within 5 km of a fishing vessel each day during the 70 days of the incubation period (see supplementary text for calculation details). In addition, assuming that the probability of interaction is proportional to the fishing effort, this number might rise to 20-25 individuals per day in winter (June-September), when sooty albatrosses from the Crozet Islands spend more time in sub-tropical waters (Delord et al. 2013). Although these estimates are based on rough approximations, and there is no information about the probability for a sooty albatross to die from bycatch when within 5km of a fishing vessel, these numbers suggest that even a low individual risk may have a significant impact on such a small population. Indeed, for a biennially-breeding species, mortalities from all sources (including bycatch) should not exceed 0.015 times the number of breeding pairs to maintain a population viable, which corresponds to 6.7 individuals per year for the sooty albatross population from Crozet (Dillingham & Fletcher 2011). The unusually low adult survival observed for this population (Rolland et al. 2010, Barbraud unpublished data), the negative relationship between adult survival and tuna longline fishing effort (Rolland et al. 2010) and the absence of population recovery since the sharp collapse observed in the 1980s suggest that individual mortality is still exceeding this rate. Further work will be needed to determine if bycatch has a substantial effect on the population trend, especially by understanding the proportion of illegal fishing vessels encountered throughout the annual cycle, as Crozet albatross populations might be more exposed to this threat (Corbeau et al. 2021b).

### Foraging behaviour and attraction to vessels

Both onboard boat observations and tracking data have demonstrated substantial inter-species variations in occurrence and interaction behaviour with fishing vessels (Hudson & Furness 1989, Cherel et al. 1996, Collet et al. 2017a, Corbeau et al. 2021a). Studies of albatross species strongly attracted to fishing vessels, such as wandering and black-browed albatrosses (*Thalassarche melanophrys*), demonstrated that individuals exhibit attractive responses by clearly changing their trajectory towards the boat when detected (Collet et al. 2015, 2017a). In the present study, there was no evidence that sooty albatrosses changed flight direction, or speed, when they were within 30 km of a vessel. The distance of 30 km is interpreted as the maximum distance at which an albatross could visually detect a vessel, although this could be dependent on the flight height of the individual, and therefore be species-specific (Collet et al. 2017a). However, the absence of any trajectory change when closer than 30 km for the sooty albatross suggests that the obvious difference in attraction towards boats compared to other species does not originate from a lower detection capacity.

When actively interacting with a boat, the behaviour of attending seabirds is similar to intense search events, with a lower speed and higher sinuosity (Torres et al. 2011, Bodey et al. 2014). Birds attracted to boats usually stay within a close range for at least a couple of hours (Collet et al. 2017b), and spend more time on the water, which is interpreted as feeding attempts (Collet et al. 2015). Sooty albatrosses in the present study, as well as in Banda et al. (2023), did not exhibit obvious signs of attraction behaviour towards boats (fishing vessels or not). In both studies, individuals remained less than 20 minutes within 5 km of the boats, did not stop, and appeared to just fly past. Although not all seabirds necessarily interact with boats that they encounter (Torres et al. 2013, Sugishita et al. 2015, Collet et al. 2017b), these results tend to confirm that sooty albatrosses are less inclined to interact with fishing vessels than other species (Griffiths 1982).

The optimal foraging theory predicts that the prey preference, the aggressiveness of individuals and their dominance rank within a seabird aggregation, could affect the decision to join or not a foraging patch generated by a fishing vessel (Giraldeau & Caraco 2000, Stephens & Krebs 1986). Therefore, the apparent low interest of sooty albatrosses towards boats could arise from competitive exclusion. The larger and/or more aggressive individuals, such as wandering albatrosses, would prevent the smaller sooty albatross from joining seabird aggregations (Weimerskirch et al. 1993, González-Solís et al. 2000). As much as this behaviour could be an inherent trait of the species, this also could have been a result of a recent human-induced evolutionary mechanism. Indeed, the sharp decrease of the sooty albatross population in the 1980s at Ile de la Possession coincides with the development of longline fisheries, and bycatch could have acted as a harvesting pressure whereby individuals more attracted to fishing vessels or more dominant in seabird aggregations were removed from the population. Such selective mortality would have had an effect on the interaction behaviour at the population level similar to what was theorised for the wandering albatross (Barbraud et al. 2013, Tuck et al. 2015).

However, even if the sooty albatross is or has become a less dominant species, or has a lower appetite for the food made available by fishing vessels, bycatch records confirm that this species is still attracted, even in a low proportion (Huang & Liu 2010). A previous study of interactions between longline vessels and seabirds recorded that sooty albatrosses were found to interact in 4% of the observations (Cherel et al. 1996). The fact that this species might only interact with fishing vessels occasionally could suggest that this behaviour is hardly detectable by short-term tracking surveys. Indeed, combining the present study and Banda et al. (2023), only 31 individuals were tracked, covering a few weeks at the end of the incubation period. Late November and early December also correspond to the period with the lowest fishing effort, suggesting that the absence of clear interaction behaviour could be an artefact of a low interaction rate, a low sample size, and few fishing vessels present in the area during the study period. In addition, attraction towards fishing vessels can vary seasonally, independently of the boat density. Factors such as low food availability and/or high energy requirements can influence seabirds by stimulating temporary high-risk behaviour (Bateson 2002, Clark et al. 2020). Therefore, the information collected about sooty albatross attraction towards fishing vessels during the incubation period may conceal a higher bycatch risk during energetically demanding periods, such as chick-rearing, moulting or pre-breeding periods.

### Conclusion and perspective

Feedback from the South Georgia longline fishery indicates that strict implementation of regulation measures, such as night-setting and line-weighting, reduced seabird mortality to negligible levels (Collins et al. 2021). However, without 100% observer coverage enforced, the level of compliance and bycatch rates remain uncertain, in particular for fisheries operating outside EEZs where they are not legally obligated to report bycatch rates. Sooty albatrosses from the southern Indian Ocean forage mostly in the High Seas (present study, Delord et al. 2013, Schoombie et al. 2017, Heerah et al. 2019, Banda et al. 2023), and although the present study may suggest that this albatross species does not exhibit a strong attraction behaviour towards fishing vessels, some individuals are still victim of bycatch (Gales et al. 1998, Huang & Liu 2010, unpublished French Southern Breeding Seabird Survey database). The absence of long-term population recovery supports the suggestion that even a low risk of bycatch at the individual level may have a non-negligible impact on this small population. In this context, it is crucial to determine if the interaction rate and attraction behaviour of sooty albatrosses change seasonally in accordance with food availability, energy requirements and fishing effort. The present study contributes to a better understanding of the sooty albatross at-sea behaviour and allows to assess the risk of bycatch during the incubation period, nevertheless, the species conservation status involves new data collection strategies. In particular, overcoming the challenges of investigating the behaviour of such species during the non-breeding period (no current accurate location data) appears to be a priority to provide a reliable assessment of the actual bycatch risk that they are facing.

## Acknowledgements

This study was funded by an Agreement on the Conservation of Albatrosses and Petrels (ACAP) Small Grant (Grant number 2020-15 to Christophe Barbraud) and by Réserve Naturelle Nationale des Terres Australes Françaises (Célia Lesage). The fieldwork in Crozet was funded and logistically supported by IPEV (project 109 OrnithoEco). We warmly thank Alexandre Vong, Jeanne Abbou, Célia Lesage (RN TAF), Nicolas Croizet (RN TAF), Clément Jaunas, Thomas Bobillier for their help in the field, and Henri Weimerskirch for his advice during the preparation of the field work. We thank Dominique Filippi from Sextant Technology for his help with the start-up of XAIS loggers. This study is part of the long-term Studies in Ecology and Evolution (SEE-Life) program of the CNRS. We would also like to thank Daphnis De Pooter (CCAMLR) and Pierre Peries (SIOFA) for assisting with the provision of the fishery data, and all data owners for their permission for the use of the data. The field procedures and manipulations on Crozet were approved by Comité National de la Protection de la Nature and by the Préfet of Terres Australes et Antarctiques Françaises (permit number 2022-70 and 2022-84).

---

**Supplementary Figure 1:**
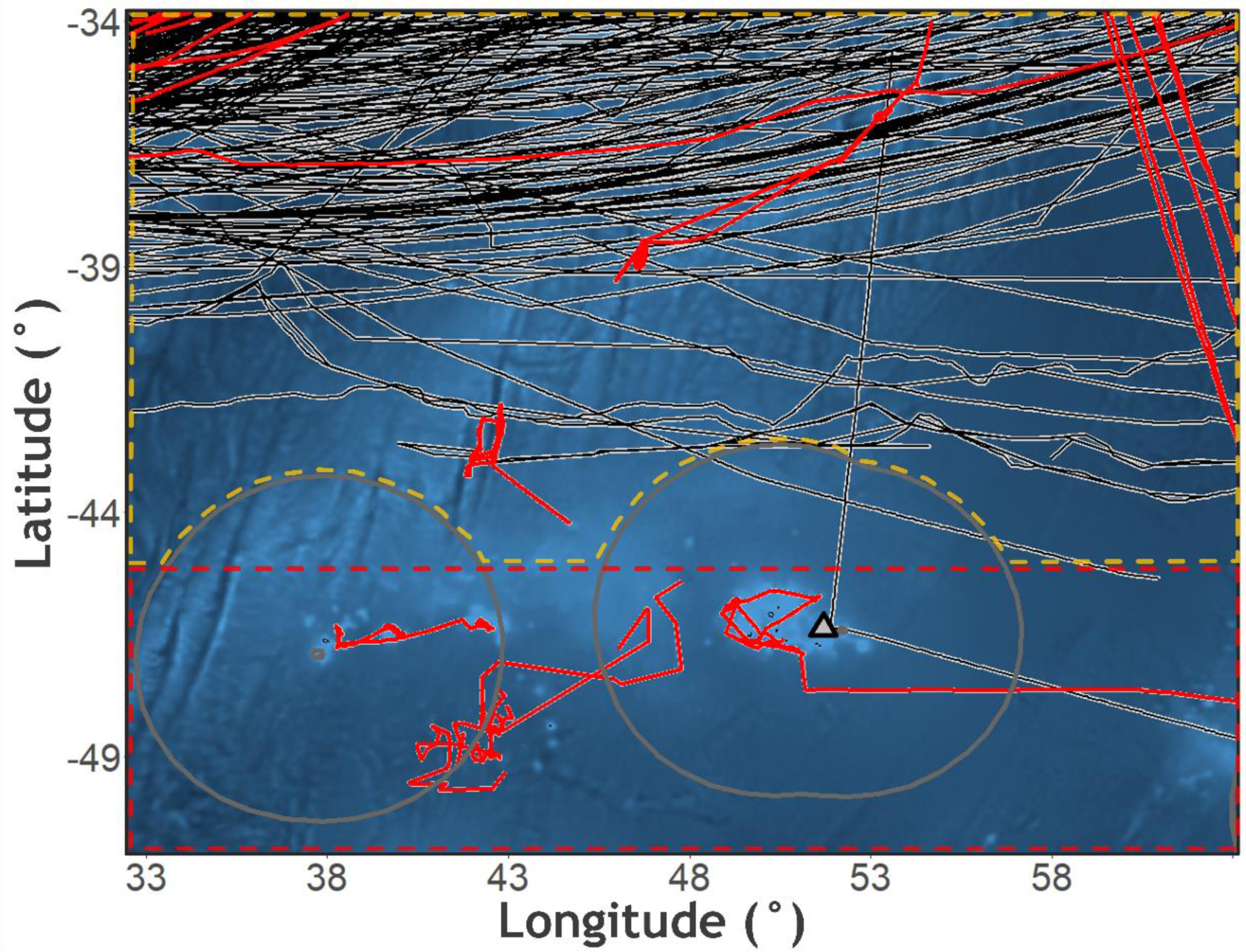
Tracks of vessels during the study period obtained from Global Fishing Watch. Tracks in red correspond to fishing vessels. The circular grey shapes correspond the Exclusive Economic Zones (EEZs) of Crozet Islands (grey triangle) and Marion and Prince Edward Islands. The dotted lines correspond to the SIOFA (orange) and the CAMMLR (red) areas.

## Supplementary text: Estimation of the number of individuals interacting with fishing vessels per day

In the present study, there was one interaction with a fishing vessel during the 11 foraging trips recorded in incubation (11 individuals) for a total of 105 days at-sea. Therefore, for this dataset, the number of interaction per day and per individual was (1/11)/105 = 0.0009. For the whole Crozet population, the number of breeding pairs was estimated to 444 in 2022-2023 (extrapolation obtained from the population monitored annually at Ile de la Possession, see Methods). For biennially-breeding albatrosses, the estimated number of individuals per breeding pair is 7.3 (conservative estimate from Dillingham & Fletcher 2011), which corresponds to 3,241 sooty albatross individuals for the whole Crozet population. With 0.0009 interaction with fishing vessels per day and per individual, the total number of individuals interacting per day was estimated to 2.9 for the whole population during the incubation period. If we consider that the interaction rate is proportional to the fishing effort, the fact that the fishing effort is 8 times higher in August than December in the study area brings an estimate of about 23 individuals interacting per day. We recommend to take these estimates with caution as they are calculated from a small dataset. This approach is used here to demonstrate that a small interaction rate can be potentially impactful at the population level.

## Notes

### Competing Interest Statement

The authors have declared no competing interest.

